# Epigallocatechin-3-gallate yield in different temperature gradients in green tea (*Camellia sinensis*) brewing

**DOI:** 10.1101/396234

**Authors:** Meng Hsuen Hsieh, Meng Ju Hsieh, Chi-Rei Wu, Wen-Huang Peng, Ming-Tsuen Hsieh, Chia-Chang Hsieh

**Affiliations:** Department of Electrical Engineering and Computer Science, University of California, Berkeley, Berkeley, California, US; Department of Medicine, Poznan University of Medical Science, Poznan, Poland; Institute of Chinese Pharmaceutical Sciences, Department of Pharmacy, China Medical University, Taichung, Taiwan, R.O.C.; Department of Pediatrics, China Medical University Children’s Hospital, China Medical University, Taichung, Taiwan, R.O.C.

**Keywords:** EGCG, green tea, antioxidant, HPLC

## Abstract

**Introduction:** Epigallocatechin-3-gallate (EGCG) is a chemical catechin, a natural organic compound found in green teas with strong antioxidative effects. EGCG degrades or epimerizes according to temperature, fluctuating its concentration in green tea (*Camellia sinensis*). This study is conducted to determine the specified correlation between EGCG and tea temperature, and to conclude with the optimal temperature for EGCG yield.

**Methods:** EGCG concentrations in different solutions of green tea are analyzed using a high-performance liquid chromatography (HPLC), with a diode array detector (DAD). The solutions are created from green tea brewed in water from 20°C to 100°C at increments of 20°C and undergo an ultrasonic bath of 30 minutes before being analyzed.

**Results:** There is a discernible difference between EGCG concentrations in all temperatures. At 20, 40, 60, 80 and 100°C, the concentrations are 6.18 μg/mL, 32.37 μg/mL, 57.36 μg/mL, 36.13 μg/mL, and 44.85 μg/mL, respectively. EGCG concentration maximizes at 60°C. The lowest EGCG concentration yield is at 20°C.

**Conclusion:** The results of our experiments lead us to recommend hot brewing over cold brewing for green tea if one wishes to maximize the potential of the effects of EGCG due to its higher concentration.

## INTRODUCTION

Green tea (*Camellia sinensis*), one of the most common beverages in the world, is usually brewed in cold or hot water. The beverage has been claimed to have an array of health benefits including anticancer and antioxidant abilities ^[1] [2]^. The agents responsible for health improvement from tea are known as catechins. There are four major types of catechins in tea, epigallocatechin-3-gallate (EGCG), gallocatechin gallate (GCG) epigallocatechin (EGC), epicatechin gallate (ECG), and epicatechin (EC), with EGCG being the most abundant among all catechin types ^[3]^. As EGCG contains eight hydroxyl (OH^-^) groups, it can remove multiple free radicals which allows it to become one of the more potent antioxidants ^[3] [2]^, additional researches show that the chemical has anti-cancer ^[1]^, anti-inflammatory effects and can improve learning and memory retention skills ^[4]^.

The stability of EGCG changes under different conditions: in higher temperatures, EGCG concentration increases due to the epimerization of GCG to EGCG, while in lower temperatures, EGCG epimerizes into GCG; both reactions change EGCG concentrations in solution. ^[5] [6]^ As such, the two principle brewing methods (cold and hot brewing) would affect the yield of EGCG, determining the healthiness of the tea preparation. A study claimed that among commonly consumed tea, green tea possesses one of the highest amounts of EGCG, yet there is no investigation into the relationship between temperature and EGCG yield in green tea solutions. ^[6]^ This study is conducted to determine the specified correlation between EGCG and tea temperature, and to conclude with the optimal temperature for EGCG yield.

## METHODS and MATERIALS

### Preparing the green tea solutions

One gram of green tea leaves was measured and macerated in five different temperatures of 10 mL water, from 20°C to 100°C at increments of 20°C. The solutions then were placed in an ultrasonic cleaner (3510R-DTH Bransonic, CT, USA), at 42 kHz for thirty minutes to brew. The solutions were lastly filtered with 0.45 μm filter papers for HPLC analysis.

### Ultrasonication process

As ultrasonication is known to break down plant cells such as tea, ^[7]^ it could rupture the cells of the leaves, allowing it to release its effluences, ^[8]^ thus amplifying EGCG yield. The amplification of EGCG concentration by ultrasonication will aid to examine the difference between different brewing temperatures by amplifying the differences among samples. ^[9]^

#### The usage of the high-performance liquid chromatography diode array detector (HPLC-DAD) machine

An HPLC-DAD machine (SPD-M10A, Shimadzu, Kyoto, Japan) was used to analyze the concentration of EGCG in the solutions. EGCG has a retention time of 30 minutes and an ultraviolet spectrum at around 280 nm ^[6]^. The HPLC mobile phase used for the machine is 100% pure methanol alongside 0.2% of acetic acid. Standard HPLC procedures by Ju, et al. apply. ^[10]^ The trials were measured three times to reduce random error.

#### Complete sample solutions and standard EGCG solutions

Since the units of measure for EGCG amount presented in the HPLC were shown as milli absorbance units (mAU), an EGCG standard solution was required for a conversion from mAU to the concentration of an EGCG solution in μg/mL. Five differently concentrated standard samples of 10 μg/mL, 20 μg/mL, 40 μg/mL, 60 μg/mL, 80 μg/mL, and 100 μg/mL were created. The standard solutions were then graphed and an equation derived from the graph to interpolate the concentration of EGCG in the tea specimens.

## RESULTS

### Stock solution data and equation

Results of the evaluation of the EGCG concentration of the standard solutions are measured in milli absorbance units (mAU) in Table 1. This set of data comes from the standard solutions. The data in Table 1 forms a positive linear correlation between the concentration of the standard solutions and the milli absorbance units. The relationship between the concentration and the measured data is *f(x)=291834.33 x-1597929.78*, where *f(x)* is the measured data in mAU, and *x* is the EGCG concentration in μg/mL.

**Table 1:**
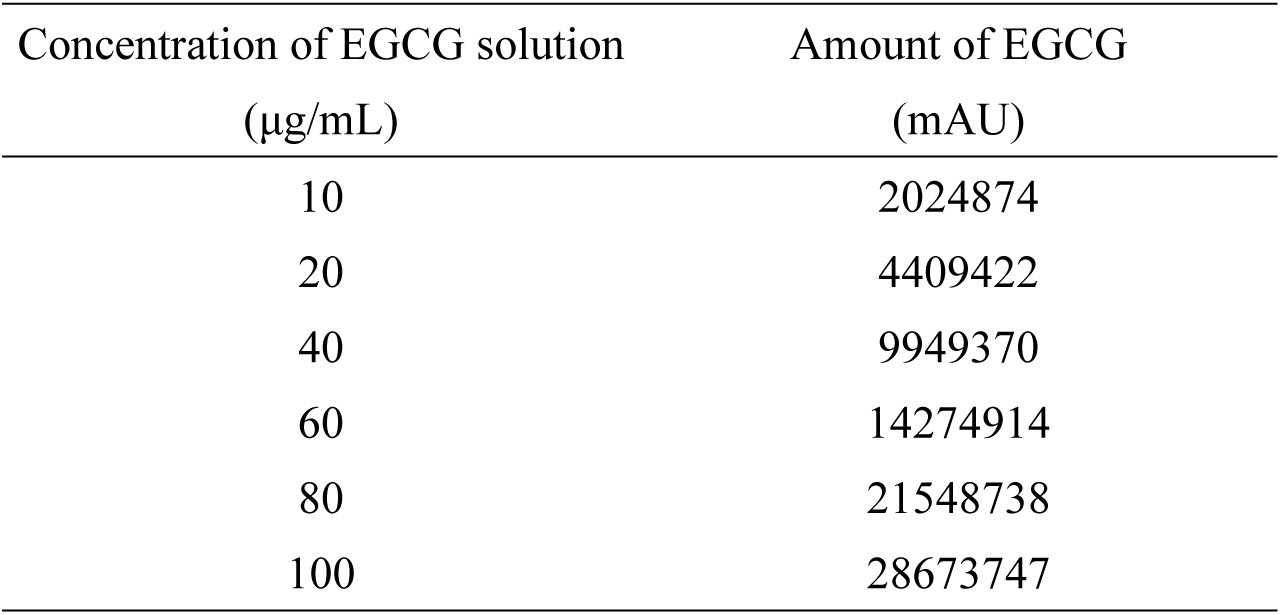
Data of the standard solutions

| Concentration of EGCG solution<br>($\mu\text{g/mL}$) | Amount of EGCG<br>(mAU) |
| --- | --- |
| 10 | 2024874 |
| 20 | 4409422 |
| 40 | 9949370 |
| 60 | 14274914 |
| 80 | 21548738 |
| 100 | 28673747 |

The data of the standard solutions is used to calculate the EGCG amount in μg/mL in the sample solutions. The graph of table 1 is located at figure 1.

**Fig 1:**
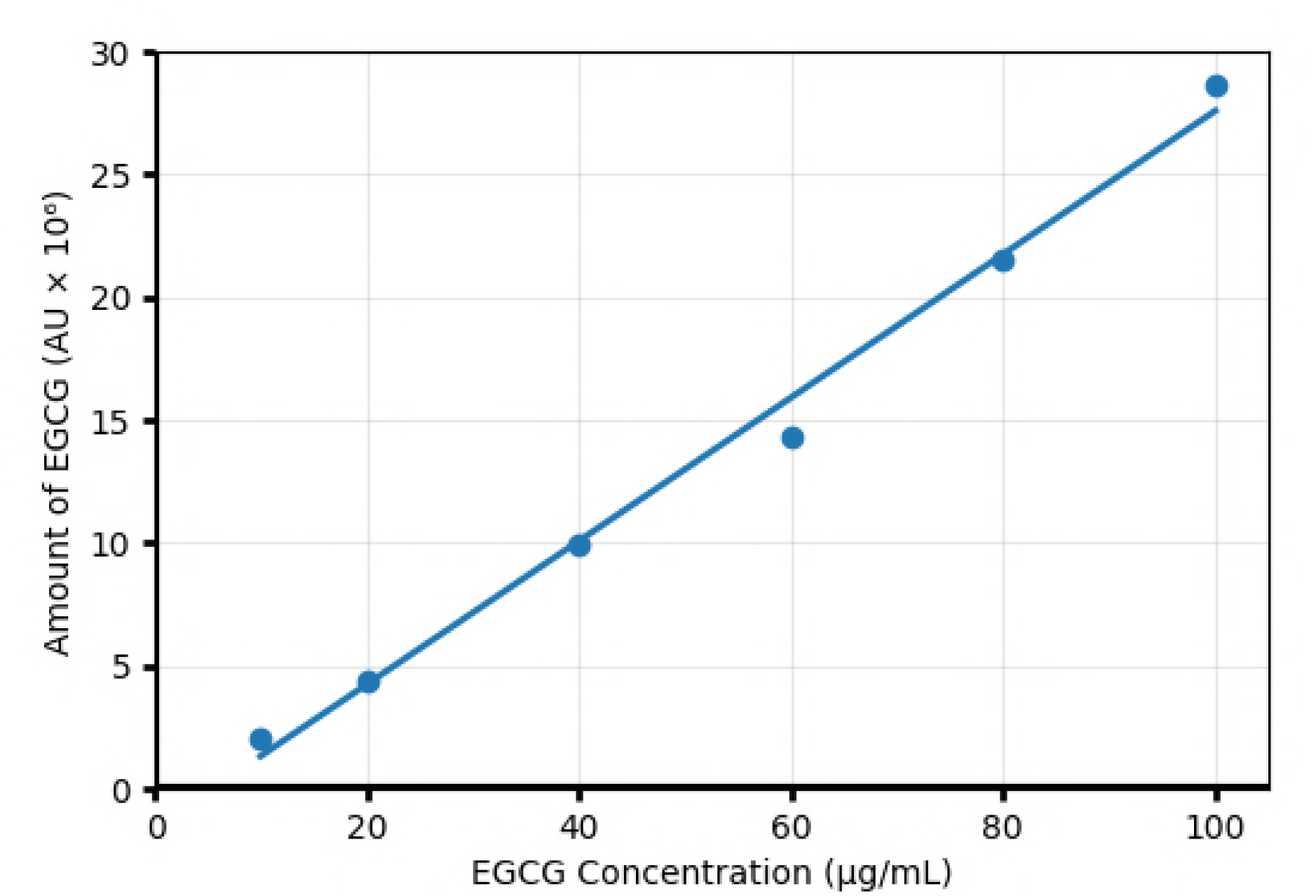
Analysis of EGCG solutions by HPLC

### Sample solution data

A total of three individual trials are conducted for each sample solution and their averages shown. The results are presented in Table 2 for the sample solutions in μg/mL as converted from the stock solution results in Table 1. The data is expressed as mean ± two standard deviations (SD). A graphical representation of table 2 is in Figure 2. EGCG concentration in the solutions slowly increases from 20°C (6.14 μg/mL) and peaks at 60°C (57.36 μg/mL). The concentration decreases slightly at 80°C (36.13 μg/mL) and increases again for the 100°C sample (44.85 μg/mL).

**Table 2:** Data of the sample solutions after analysis by the HPLC machine

| Solution Temperature<br>(°C) | EGCG amount (µg/mL) |  |  |  |
| --- | --- | --- | --- | --- |
|  | Test 1 | Test 2 | Test 3 | Average |
| 20 | 6.14 | 6.18 | 6.22 | 6.18 |
| 40 | 33.03 | 31.46 | 32.63 | 32.37 |
| 60 | 52.78 | 59.30 | 60.00 | 57.36 |
| 80 | 33.50 | 37.26 | 37.61 | 36.13 |
| 100 | 43.63 | 45.17 | 45.76 | 44.85 |

**Fig 2:**
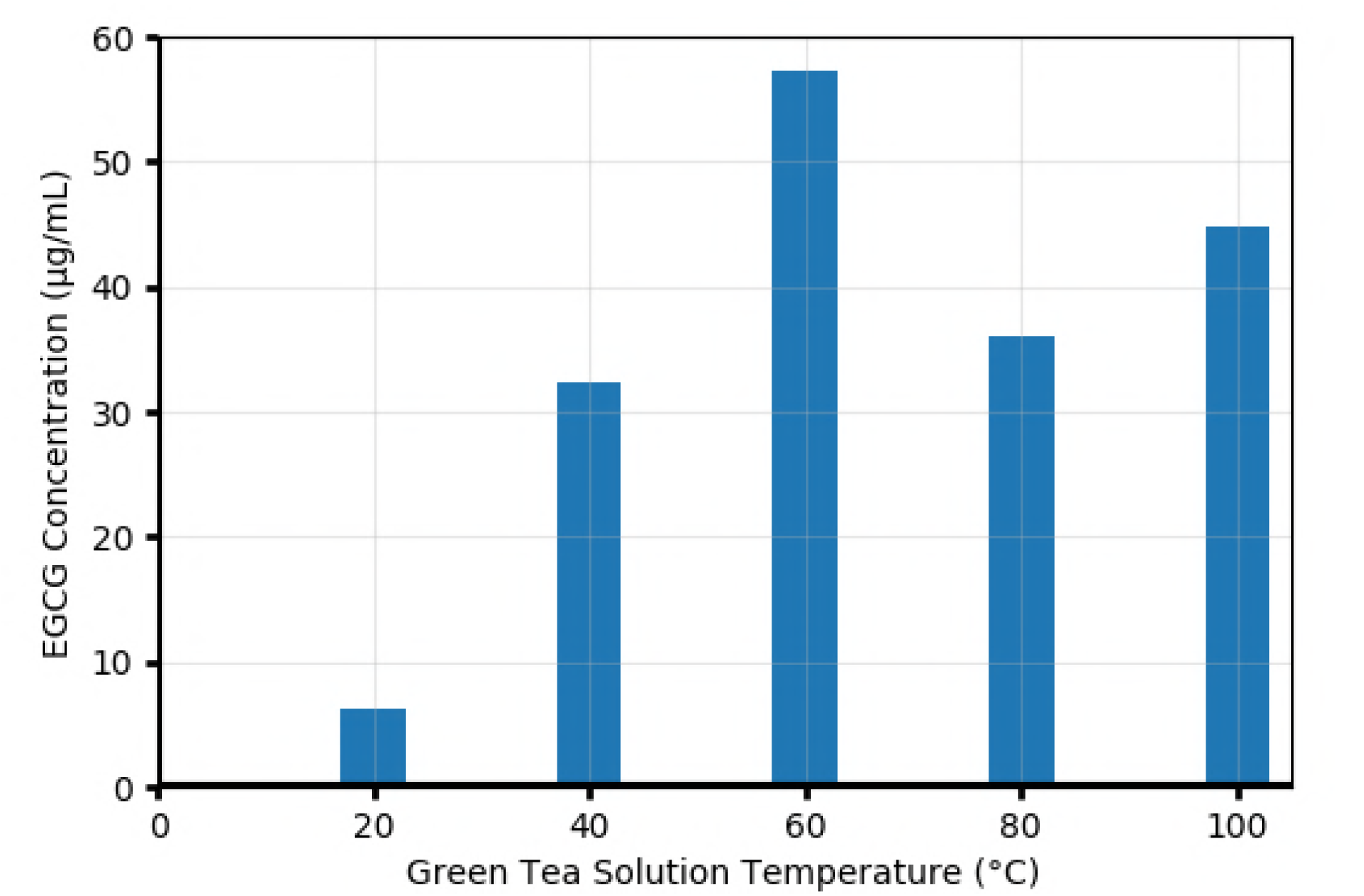
EGCG yield in relation to temperature

## DISCUSSION

The results of our study show that the EGCG concentration varies among different temperatures, with a clear peak at around 60°C at 57.36 μg/mL. Since no external changes are done to the solutions except for controlling the temperature, the fluctuations in concentration would come from changes within the solution. EGCG epimerizes to GCG and vice versa at different temperatures. ^[11]^ Below 44°C, the EGCG yield decreases as it epimerizes into GCG; above 44°C, the rate of epimerization of GCG into EGCG overtakes the epimerization of EGCG into GCG, thus increasing EGCG concentration after 44°C. At 98°C, the rate of GCG epimerization into EGCG increases relatively significantly ^[5]^. The results of our experiments corroborate these findings well; there is a notable increase in EGCG yield starting from 40°C, peaking at 60°C, and again increasing at the 100°C mark.

The reason for the decrease in EGCG concentration at 80°C (36.13 μg/mL) from the peak at 60°C (57.36 μg/mL) is not known. Other studies have noted there to be a turning point in EGCG/GCG epimerization and degradation at 82°C ^[5]^. The results of this study show the same observation, but does not provide a suitable explanation. Further studies can be performed to examine the phenomenon.

On a strictly EGCG-based recommendation, a 60°C brewed green tea would provide the most potent antioxidant effects due to it possessing the highest concentration of EGCG. Despite that, controlling water temperatures to remain at 60°C may be a difficulty for quotidian tea brewing with hot water boilers. As such, due to the relatively low discrepancy between the 60°C and 100°C data points (13.01 μg/mL difference), for practical purposes, the recommended temperature for green tea brewing to optimize EGCG concentration would be at a boiling 100°C. Cold brewing (25°C), in this case, would not be recommended over hot brewing, since any temperature above 20°C yielded a higher EGCG concentration.

Ultrasonication processes have been investigated previously in other studies for catechin yields in green tea solutions, noting that ultrasonication does increase catechin yields. ^[12]^ Though this study did not conduct any procedures to corroborate these findings, the discernible differences seen among the different tea samples suggest that ultrasonication can be purposeful in future experiments seeking to investigate catechin concentration yields in different environments.

The limitations of this experiment lie in the large temperature increments (20°C apart). The optimal temperature for EGCG concentration maximization can only be approximated to be around 60°C; the true value can be anywhere between 40°C and 80°C. Though the exact temperature cannot be determined from this study, it was able to conclude that hot brewing does provide more potent EGCG benefits over cold brewing.

## CONCLUSION

The results of our experiment show that there is a discernible difference in the concentrations of the chemical makeup of green tea in different temperature solutions. EGCG concentration is significantly higher in warmer brewing than in colder brewing. This leads us to recommend hot brewing over cold brewing for green tea if one wishes to maximize the potential of the effects of EGCG. Additionally, ultrasonication provides both research potential for future experimental designs and commercial potential in its ability to increase EGCG concentrations.

## ACKNOWLEDGEMENTS

We would like to thank Dr. Chi-rei Wu for allowing us to conduct the experiment under his guidance in his laboratory at China Medical University in Taichung, Taiwan.

